# Unveiling vertebrate biodiversity in arid and semi-arid terrestrial ecosystems through eDNA metabarcoding at savanna waterholes

**DOI:** 10.1101/2025.10.06.680638

**Authors:** Tamara Schenekar, Janine Baxter, Irmgard Sedlmayr, Julia Gladitsch, Sibusiso Mahlangu, Monica Mwale

## Abstract

Applying environmental DNA (eDNA) metabarcoding to samples from waterholes and their surroundings offers a promising approach for monitoring terrestrial vertebrates in semi-arid and arid ecosystems, such as the southern African savannas. However, minimal guidance exists on key sampling design parameters for terrestrial ecosystems, which can significantly influence species detection. This study investigated the effects of sampled substrate, sampling season, and metabarcoding primer pair on species richness and taxonomic group detection in terrestrial vertebrates, with a focus on mammals, using eDNA samples from waterholes in Botsalano Game Reserve, South Africa. A total of 725 eDNA samples were collected from 94 sampling events across wet and dry seasons, detecting 95 species (45 birds, 42 mammals, 4 amphibians, 3 reptiles, and 1 fish). Sediment samples provided more reliable detection of abundant taxa, whereas water samples had higher detection frequencies of rare taxa. A mixed sampling approach yielded the highest species richness. Sampling during the wet season yielded higher species richness overall, while more mammal species were detected from dry season sampling. Overlap in species detection between the two metabarcoding primers tested was low (47%). We formulate recommendations for future eDNA metabarcoding study designs in similar systems, including remote sampling logistics and discuss potential sources of false positives in eDNA metabarcoding, including (1) secondary eDNA input, (2) incomplete genetic reference databases, and (3) the low genetic resolution of metabarcoding markers.

## Introduction

The use of environmental DNA (eDNA), defined as extraorganismal DNA shed into the environment and extractable from environmental samples (Taberlet et al., 2012), has proven to be an effective approach in enhancing biodiversity assessments for management applications such as ecosystem health monitoring (Aylagas et al., 2014; Ruppert et al., 2019; Suren et al., 2024; Yang & Zhang, 2020), conservation planning (Cristescu & Hebert, 2018; Thomsen & Willerslev, 2015) or environmental impact assessments (Allan et al., 2023; Laroche et al., 2018).Hereby, eDNA metabarcoding has demonstrated high cost- and time-efficiency in the simultaneous detection of multiple species (Carvalho et al., 2022; Ji et al., 2013; Svenningsen et al., 2022; Valentini et al., 2016) often outperforming or complementing traditional survey methods (Fediajevaite et al., 2021). To date, there has been less focus on the development of this approach in terrestrial ecosystems compared to aquatic environments like rivers, lakes or marine systems (van der Heyde et al., 2022). This is partly due to historical reasons (early eDNA studies on macroorganisms targeting aquatic systems), partly driven by conservation and monitoring needs, but also due to practical factors such as the ease of concentrating eDNA from water through filtration and the relative homogeneity of water, which facilitates uniform sampling. In contrast, terrestrial systems exhibit a far more patchy distribution of eDNA across and within substrates, such as soil, water and plant surfaces, due to their greater heterogeneity on smaller spatial scales (Cowgill et al., 2025; Grosberg et al., 2012). This makes sampling design even more critical for terrestrial studies. To date, the most frequently sampled substrate for terrestrial systems is soil, followed by ingested or fecal material and water (Cowgill et al., 2025; van der Heyde et al., 2022). However, the optimal sampling substrate may vary depending on the target taxa (van der Heyde et al., 2020) and, although rarely employed, combining multiple substrates often increases detected species richness (Cowgill et al., 2025; van der Heyde et al., 2022).

The temporal aspect of sampling is equally crucial and has also been shown to influence eDNA-based species detection (Buxton et al., 2018; Djurhuus et al., 2020; Johnson et al., 2021). This is either due to different physical conditions of the environment across seasons, affecting eDNA persistence, or different activity patterns of the target organisms, affecting eDNA deposition (Buxton et al., 2017; Salter, 2018). However, the majority of eDNA-based studies monitoring ecological restoration in terrestrial environments, use only a single sampling period (van der Heyde et al., 2022)

The choice of the metabarcoding primer pair is a third key parameter in any eDNA metabarcoding analysis. They define the taxonomic target group of the eDNA metabarcoding assay and have been shown to affect species detection, even when targeting similar taxonomic groups (Schenekar et al., 2020; Zhang et al., 2020). Multiple universal primer pairs are increasingly used to maximize species detection, despite higher laboratory costs (Elbrecht et al., 2017; van der Heyde et al., 2022).

The biomes of the African continent still remain among the least explored systems concerning eDNA dynamics and application (Belle et al., 2019; Cowgill et al., 2025; Schenekar, 2023; von der Heyden, 2022). However, their exceptional biodiversity and persisting anthropogenic threats (Archer et al., 2021; Myers et al., 2000), demand efficient and cost-effective monitoring techniques to inform conservation efforts and guide management policies. In semi-arid to arid systems like savannas, waterholes (both natural and artificial) serve as essential aggregation points for wildlife seeking scarce water resources, making these critical locations for monitoring and management efforts (Redfern et al., 2005; Sutherland et al., 2018). Few studies have started to explore the applicability of waterhole-borne eDNA to monitor pathogen (Alfano et al., 2021; Farrell et al., 2019) or mammal (Farrell et al., 2022; Li et al., 2023; Mas-Carrió et al., 2022) biodiversity in Africa. Despite these studies revealing the high potential of waterhole-borne eDNA to monitor terrestrial wildlife, a systematic evaluation of the variability and reliability of eDNA metabarcoding while testing for multiple workflows is lacking.

This study examines how various sampling design and workflow parameters affect eDNA metabarcoding results for monitoring terrestrial vertebrates in savanna systems using samples collected from waterholes and their immediate surroundings. This was conducted through two experimental setups. The first experiment aimed at understanding the primary factors such as substrate, season and primer affecting the detection of terrestrial vertebrate species. The second experiment evaluated the potential use of soil samples from wildlife trails near waterholes as an alternative to direct sampling of water or sediment from waterholes.

## Material & Methods

### Study site

The study was conducted in Botsalano Game Reserve, a 6,000-ha game reserve situated in North West Province, South Africa (Fig. 1). The prevailing climate is semi-arid with hot, rainy summers (October to April) and cold, dry winters (May to September). Sampling was conducted during two seasons in 2023: at the end of the wet season (March 10^th^ to April 23^rd^, 2023) and at the end of the dry season (September 9^th^ to October 23^rd^, 2023, Fig. S1).

**Fig. 1.**
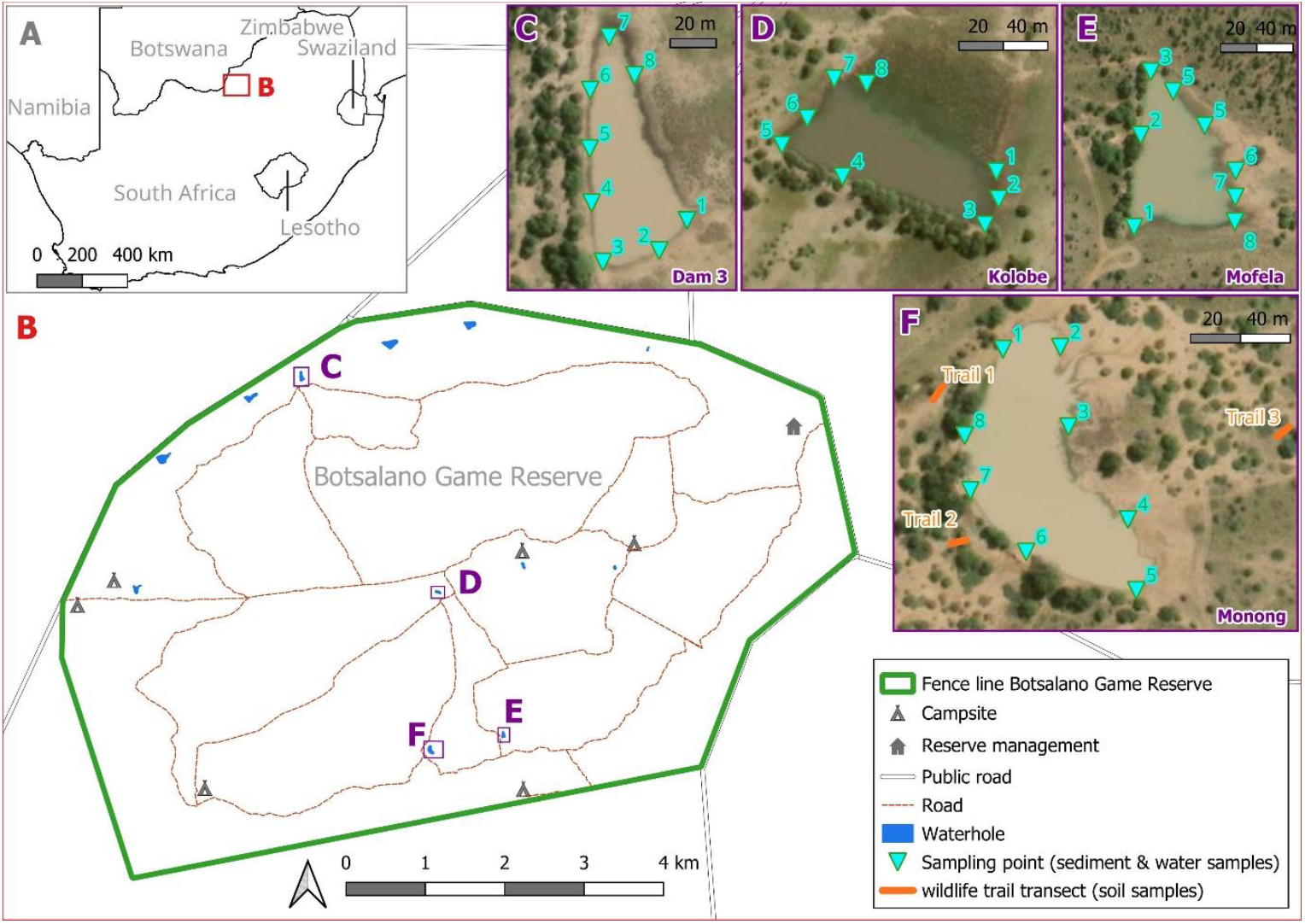
Location of Botsalano Game Reserve (A) as well as the sampled waterholes and wildlife trails within the reserve (B). Approximate sampling points at the each of the waterholes are shown in (C-F). Satellite images: ESRI (2024)

Botsalano Game Reserve lies within the savanna biome in the highveld of South Africa, encompassing umbrella thorn savanna and woodland as well as back thorn scrub (Morris, 2022). Within the reserve, dams have been constructed along ephemeral streams to hold water during the dry season, resulting in semi-natural waterholes. The ephemeral streams fall dry soon after the main rainfalls (November-February), so that all waterholes represent independent water bodies. The reserve does not conduct regular game counts for species inventory, although Morris (2022) lists more than 100 bird species and over 20 larger-bodied mammals (Table S1).

Three waterholes in the reserve were selected during each season based on accessibility, spatial distribution, and anticipated water levels during the wet and dry seasons. Although the initial sampling design intended to use the same waterholes across both seasons, one waterhole (Dam 3) dried up at the start of the dry season and was replaced with an alternative waterhole (Monong) during sampling. Consequently, the waterholes sampled during the wet season were Kolobe, Mofela and Dam 3, while Kolobe, Mofela and Monong were done during the dry season (Fig. 1).

### Sample collection & processing in the field

One waterhole or all three trails were sampled per day, with the three waterholes per season being sampled over three consecutive days, followed by a three-day break. This resulted in a six-day sampling interval for each waterhole and a total of eight sampling days per waterhole during the wet season. A ‘sampling event’ hereafter refers to the collection of a specific substrate (water, sediment, or soil) from a specific location (e.g., Waterhole ‘Mofela’ or ‘Trail 1’; see below) on a given sampling day. Due to low water levels during the dry sampling season, number of sampling events varied across waterholes (Kolobe – water 2x, sediment 5x; Mofela - water 5x, sediment 8x; Monong - water & sediment 7x). Wildlife trails were sampled in dry season only, at four sampling events.

For each sampling day at a waterhole, eight replicate samples of both water and sediment were collected. For water sampling, a DNA-free 500 mL Nalgene bottle mounted to a 2 m PVC tube was utilized to scoop up surface water approximately 1.5 m away from the shoreline (Fig. S2a). For sediment sampling, approximately 3 ml of sediments, still submerged in water, were collected directly at the shoreline and transferred into a 15 ml tube using a DNA-free spatula (Fig. S2b). During each sampling event at a waterhole, pH, conductivity, and dissolved oxygen level of the water body were measured.

For wildlife trail soil sampling, three trails around Monong waterhole, located within a 60 m radius, were selected for the placement of line transects for defining sampling points (Fig. 1F). Along each trail, a 5 m transect was established using a rope marked at 1 m intervals, creating six evenly spaced sampling points per transect from which six soil samples were collected per sampling event. Trails were sampled at 6-day intervals (all three trails on the same day) over a total of four sampling events. At each sampling point, approximately 3 mL of surface soil was collected from a 3×3 cm area using a DNA-free spatula and placed in a 15 mL tube (Fig. S3).

All samples were kept refrigerated until filtering (water samples) or preservative buffer was added (sediment and soil samples), within a maximum of five hours after sampling.

Water samples were filtered using a 250 ml Nalgene filter holder mounted onto a 1 l vacuum flask, powered by a peristaltic pump (MasterFlex peristaltic vacuum pump LS 600R) via a MasterFlex L/S 15 silicone tube and pumped at a maximum speed of 700 ml/min. Filtering was conducted until either the entire volume of 500 ml was processed or after a maximum filtration time of 10 minutes per filter. In the wet season, all samples were first prefiltered through a 20 µm Nylon filter, followed by filtration through a 3 µm MCE filter. If less than 250 ml was filtered through the 3 µm MCE filter, the remaining sample was then filtered through an 8 µm MCE filter, and this filter was used for DNA extraction and metabarcoding. In the dry season, which involved higher water turbidity, increased suspended particle load and greater filter clogging, the 20 µm Nylon filter was used directly for DNA extraction and metabarcoding. Filters were transferred to 5 ml tubes containing 2.5 ml Longmire’s solution with added sodium azide (Longmire et al., 1997). For sediment and soil samples, 6 ml of Longmire’s solution was added to each sample. All samples were stored at 8°C in the field until transport to the genetic laboratory (up to seven weeks) where they were stored at -20°C until DNA extraction. All sampling and filtering equipment (e.g., sampling bottles, PVC sampling tube, spatulas, filter holder, forceps, etc.) were cleaned between sampling events and waterholes by soaking in 30% bleach for 30 min followed by soaking in distilled water for 30 min. On each day, a field negative control was processed by filtering 150 ml of distilled water before processing the first sample of that day.

### DNA extraction, metabarcoding library preparation and sequencing

DNA extractions and setup of PCRs were conducted in a dedicated low-template DNA laboratory under two separate laminar-flows for DNA extractions and PCR setup. One extraction blank was processed with every DNA extraction batch (up to 11 samples per batch) and laminar-flow work benches were irradiated with UV light for 30 min between work sessions. DNA extractions were carried out with the DNeasy PowerSoil Pro Kit (Qiagen) using the protocol of Schenekar et al. (2024) in a final elution volume of 100 µl. For metabarcoding library preparation, two primer pairs were utilized targeting two different fragments of the mitochondrial 12S rRNA fragment: 1. The MiMammal primer set designed to amplify mammalian sequences and 2. The 12SV5 primer designed to identify vertebrate sequences (Table 1). Both primer pairs were fitted with eight-base-pair (8-bp) indices on the 5’ ends of both the forward and reverse primers, respectively, to enable sample demultiplexing using unique dual indices (UDIs). A total of four PCR (polymerase chain reaction) replicates were analysed per sample for amplification of both markers and then pooled. Each PCR consisted of 1.5 µl template DNA, 0.25 µl Q5 High-Fidelity Polymerase (2,000 U/ml, New England Biolabs), 2.5 µl Q5 Reaction Buffer, 0.2 mM dNTPs, 0.3 µM of forward and reverse primer, respectively and nuclease-free water up to final reaction volume of 12.5 µl. Cycling conditions were as follows: Initial denaturation at 98°C for 30 sec, followed by 40 cycles of denaturation at 98°C for 20 sec, annealing at 64°C (MiMammal) or 57°C (12SV5) for 30 sec, extension at 72°C for 60 sec and a final extension at 72°C for 7 min. Due to co-amplification of non-target fragments from algae and bacteria, resulting in double bands, all PCR products were excised from a 2% agarose gel and subsequently cleaned using the Monarch Spin DNA Gel Extraction Kit (New England Biolabs). Each PCR batch encompassed the amplicons of 48 samples per marker and included two controls: a PCR positive control/mock community and a PCR negative control (HPLC-purified water). The mock community contained the DNA extracts (7 ng/µl) of six non-native mammals, namely Eurasian beaver (*Castor fiber*), European hedgehog (*Erinaceus europaeus*), raccoon (*Procyon lotor*), brown bear (*Ursus arctos*), Eurasian fish otter (*Lutra lutra*) and roe deer (*Capreolus capreolus*). Field and extraction negative controls were processed in the same manner as field samples. A total of 192 amplicons (96 samples, two markers each) were pooled per library and sent to Novogene for final library preparation (using the NEBNext® Ultra™ II DNA PCR-free Library Prep Kit for Illumina, New England Biolabs) for illumina adapter incorporation and illumina sequencing. Libraries were sequenced on an illumina NovaSeq X in 150 bp paired-end read mode with a targeted output of 80 mio reads per library (417,000 reads per sample and marker).

**Table 1.** The primer pairs utilized for DNA metabarcoding. Given are the primer pair name, the sequences of forward and reverse primer, respectively, the approximate target amplicon length and the original reference of the primer pair. XXXXXXXX indicate 8-bp index sequences that were used for UDI sample demultiplexing.

| Primer pair | Primer sequences | amplicon length | Reference |
| --- | --- | --- | --- |
| 12SV5 | forward: 5'-XXXXXXXXYAGAACAGGCTCCTCTAG-3'<br>reverse: 5'-XXXXXXXXTTAGATACCCCACTATGC-3' | 98 bp | Riaz et al. (2011) |
| MiMammal | forward: 5'-XXXXXXXXGGRYTGGTHAATTTCGTGCCAGC-3'<br>reverse: 5'-XXXXXXXXCATAGTGRGGTATCTAATCYCAGTTTG-3' | 171 bp | Ushio et al. (2017) |

### Bioinformatic analysis and taxonomic classification of ASVs

Raw reads were quality-checked with FastQC (Andrews, 2010). Quality filtering was conducted with the *fastx_filter* function of *vsearch* v2.15.1 (Rognes et al., 2016) with the maximum error rate set to fastq_maxee = 1. Paired-end reads were merged using the *fastq_mergepairs* function, using the default settings and allowing for merging of staggered reads. Samples were demultiplexed by indices (1^st^ level) and the by primer pair (2^nd^ level) allowing for one error in the index and allowing for two errors in the primer sequences using *cutadapt* 4.1 (Martin, 2011). *Cutadapt* was subsequently also used for length filtering. Hereby, only reads with a minimum length of 85/155 bp and a maximum length of 115/190 bp were kept for 12SV5 and MiMammal, respectively. Unique amplicon sequence variants (ASVs) were created from the entire dataset using the *derep_fulllength* function and denoising was conducted by only retaining ASVs that had a minimum read count of five reads and a minimum alpha of 2 using the *unoise_alpha* algorithm of *vsearch*. Chimera sequences were removed from the ASV dataset using *uchime3_denovo* function of *vsearch*. Quality-filtered reads were mapped to ASVs using the *usearch_global* function with a threshold of 99% similarity. Taxonomic assignment was done using the srRNA_SINTAX database of MIDORI2 (v. GB261; Leray et al., 2022) via the *sintax* algorithm of *vsearch* and a sintax_cutoff of 0.9.

A list of previously documented species in South Africa was retrieved querying the GBIF database (GBIF.org, 2025) using the following settings: Country: South Africa, Year: 1950 and later; Basis of record: Observation & Human Observation. For detected species that did not have any occurrence records (hereafter called “non-documented species”), a species list of congeneric species that do have occurrence records based on these criteria was compiled. Additionally, a species list with the same criteria was compiled for all classifications that could only be resolved to the genus level. Both species lists were checked against the MIDORI2 database, using a custom bash script, to check which species lacked 12SrRNA records. For taxonomic assignment, if the genus or the non-documented species could be matched to a previously documented species, i.e., only one congeneric species occurring in South Africa, it was assigned to that species. If the non-native species or genus could not be unambiguously assigned to one native species, the taxon and corresponding ASVs were excluded from the analysis, with the reason for exclusion listed in Table S3. Furthermore, any detection of domestic animals, humans, or that likely stem from anthropogenic input through food or contamination (e.g., *Thunnus albacares, Salmo salar*) were discarded. The only exception was the detection on *Canis lupus*, being re-classified as *Canis mesomelas*/*Lupulella mesomelas* (black-backed jackal) as this was an abundant native species in the reserve that has only one genetic reference sequence in MIDORI2.

### Statistical analysis

Statistical analyses were performed separately for each primer pair dataset (12SV5 and MiMammal), and as a combined data set including detections from either primer pair. Species richness (number of detected species) was calculated across the full dataset and separately by location (e.g., Kolobe, Monong, Trail1), substrate (water, sediment, soil), season (dry/wet), and primers. Species richness was also assessed for all taxa combined and by taxonomic class. Euler diagrams of shared/exclusive species were created using *eulerr* (Larsson, 2024) in *R 4*.*4*.*2* (R Core Team, 2023) with *RStudio* (Posit team, 2024). For each detected species, the number of positive samples per sampling event was recorded (combined dataset only). For further analyses, species detections from all samples per sampling event were pooled (eight samples for water/sediment, six for soil). Kruskal-Wallis tests assessed richness differences by location, substrate, season, and primer. Nonmetric Multidimensional Scaling (NMDS) plots were created using *metaMDS* of *vegan* (Oksanen et al., 2025). Permutational Multivariate Analysis of Variance (PERMANOVA, 999 permutations; Bray-Curtis dissimilarity distances) was conducted using *adonis2* of *vegan* to assess the effects of location, substrate, season, primer and days since first sampling day on detected species composition. Species accumulation curves were generated with *specaccum* (*vegan)*. Species-specific detection frequency was calculated as the proportion of sampling events detecting a species (per substrate and overall). A Wilcoxon Signed-Rank Test compared detection frequencies between birds and mammals, and between sediment and water, the latter for all taxa combined as well as for birds/mammals separately. Species were categorized as ‘rare’ (overall detection frequency < 0.1) or ‘abundant’ (≥ 0.1), and detection frequencies were compared between these two groups. A hypothetical mixed-substrate dataset was created as follows: 1) For all sampling days where eight water samples and eight sediment samples were collected (N = 38), we calculated mean species richness per sampling event as well as overall species richness for a) sediment (all eight sediment samples), b) water (all eight water samples), and c) mixed-substrate samples (four randomly chosen sediment samples and four randomly chosen water samples). Kruskal-Wallis tests evaluated whether mean species richness differed among these sampling approaches. Data handling and data visualization used *dplyr* (Wickham et al., 2023), *readr* (Wickham, Hester, et al., 2024), *tidyr* (Wickham, Vaughan, et al., 2024), *ggplot2* (Wickham, 2016), *gridExtra* (Auguie, 2017), *cowplot* (Wilke, 2024) and *RColorBrewer* (Neuwirth, 2022).

## Results

A total of 725 field samples (301 water, 352 sediment and 72 soil samples) were collected in 94 sampling events, along with 38 field negative controls across both sampling seasons (Table 2, Table S4). Overall water temperature was lower and conductivity was considerably higher in dry season. The mean filtered volume of water samples collected in the dry season, was larger than those from the wet season despite higher turbidity and conductivity, due to the larger pore size of the filters used in this season.

**Table 2.**
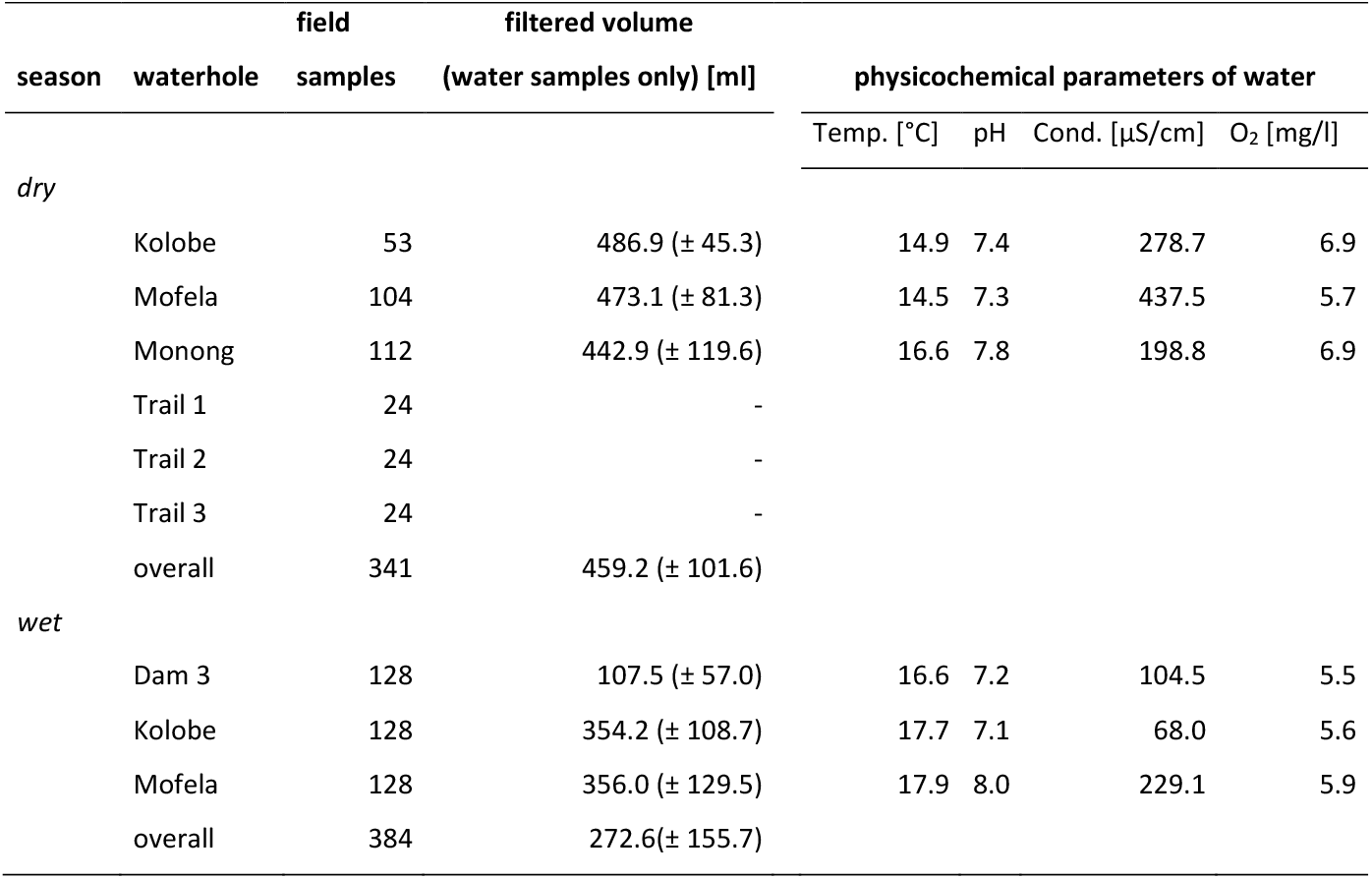
Overview of field sample numbers (field samples) for each waterhole and wildlife trail, separated by season. For water samples, the filtered volume (mean ± stdev) is given. Additionally, the mean values of the physicochemical parameters for the three water holes during the two sampling seasons are given.

A total of 878 samples (725 field samples, 38 field negative controls, 77 extraction blanks, 19 PCR positive controls, 19 PCR negative controls) were sequenced. Total raw data output was 759,743,937 reads, with 499,499,743 being retained after read quality filtering, pair merging, demultiplexing and length filtering. Dereplication identified 402,263 initial ASVs for 12SV5 and 724,237 initial ASVs for MiMammal. Of these, 2,566 and 1,683 were kept after denoising and chimera removal for 12SV5 and MiMammal, respectively. Of the retained ASVs, 354/492 could be assigned to species level and 104/178 genus level. A total of 410,190,196 reads (212,417,004 for 12SV5 and 197,773,192 for MiMammal) were assigned to an ASV at species or genus level and were used in the final dataset. Among these, field samples had on average 273,504 reads assigned for the 12SV5 dataset and 250,875 reads assigned for the MiMammal dataset, with the field and laboratory blanks having on average less than 10% of those read numbers (except for field blanks for MiMammal at 16%, Tables S5 & S6).

Initially, sequences were identified as belonging to 130 species and additional taxa belonging to 44 genera in the entire sequencing dataset. After taxonomic reassignments based on GBIF occurrences, 95 species were retained in the final dataset (Table S3, Fig. 2). The 12SV5 dataset detected 72 species while 68 species were detected with the MiMammal dataset (Table 3). The highest species richness was observed at Mofela Dam, which also had the largest number of samples and sampling events. Sediment and soil samples revealed nearly equal species richness, with 76 and 77 species detected, respectively while only 38 species were detected from soil samples. Water samples had the highest number of exclusive species (Table 3), while soil had the lowest. Overall, more species were detected during the wet season (n=78) compared to the dry season (n=70). A total of 29 species (31%) were detected in all three substrates, 53 species (56%) were detected in both seasons and 45 species (47%) were detected by both primer pairs (Fig. 3). Kruskal-Wallis tests showed significant differences in number of detected species among locations (H = 24.8, p < 0.001), substrates (H = 19.6, p < 0.001) and between primer pair datasets (H = 11.0, p < 0.001) but not between seasons (H = 0.4, p = 0.537).

**Table 3.**
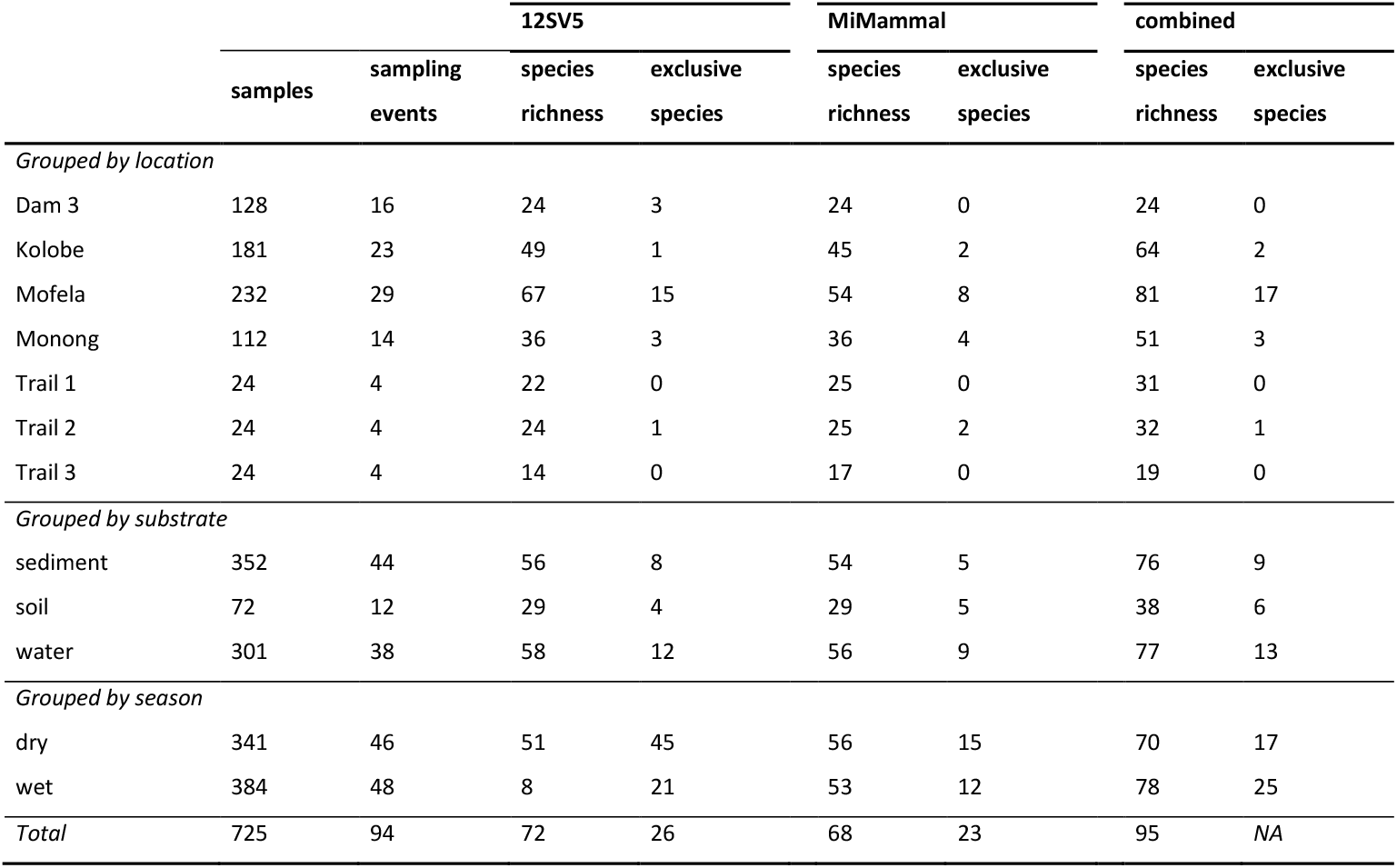
Number of species detected across different locations, substrate and seasons. For each dataset, the number of samples (samples), sampling events (sampling events), total number of detected species (species richness), and number of species that were detected only in the respective dataset (exclusive species) are shown. Data is given for 12SV5 and MiMammal datasets, as well as the combined dataset.

**Fig. 2.**
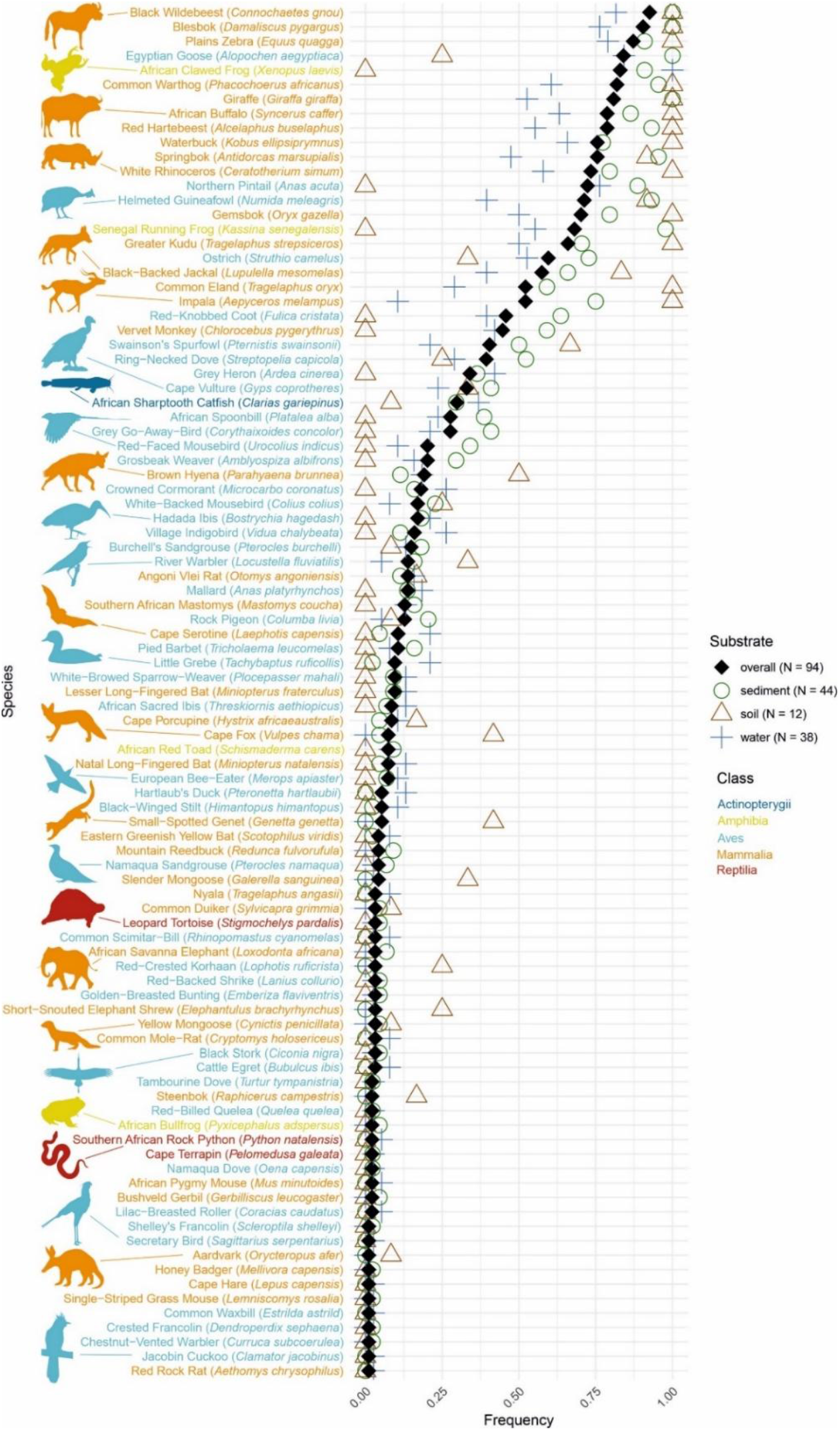
All 95 species detected by eDNA metabarcoding of sediment, soil and water samples, sorted by their overall detection frequencies in the 94 sampling events (black diamonds). Colored symbols (green circles, brown triangles and blue crosses) indicate the detection frequencies across sampling events in the respective substrate of each species.

**Fig. 3.**
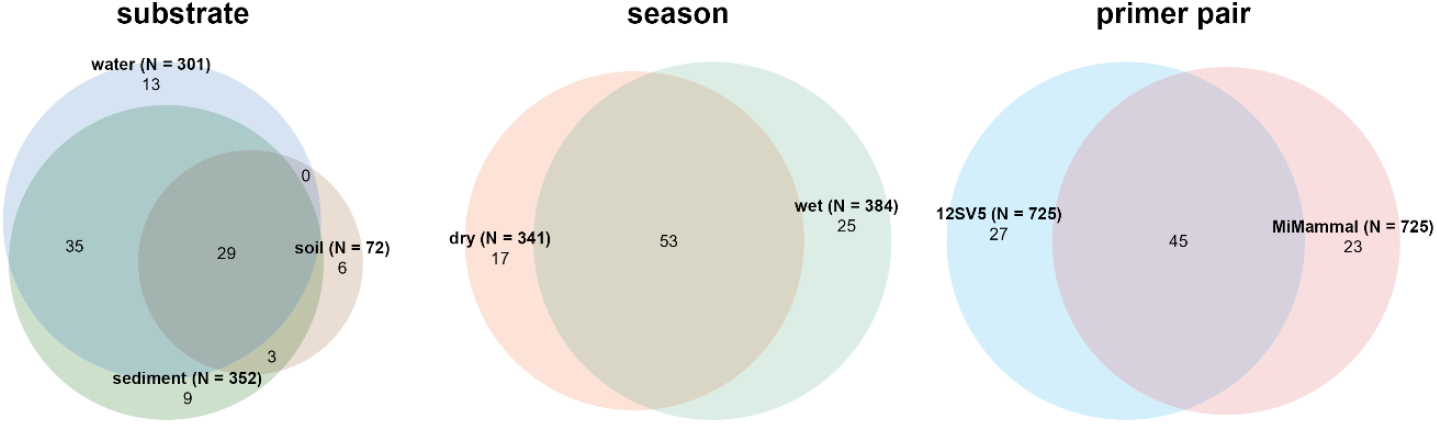
Euler diagrams showing the number of species detected across datasets for different substrates, seasons, and primer pairs. Bold labels indicate the dataset names, as well as number of samples in brackets. Regular font numbers represent the number of species unique to a dataset or shared between datasets in the respective overlaps. Diagrams for substrate and season show the dataset over both primer pairs combined.

Overall, birds were the most species-rich class (45 species), followed by mammals (42 species). The 12SV5 primer pair identified species from Actinopterygii, Amphibia, Aves, Mammalia, and Reptilia. Notably, the MiMammal primer pair detected species from all these classes, except Reptilia (Fig. 4, Table S7) and identified more mammal species (n=37) than the 12SV5 primer pair (n=29). Seasonal differences in taxonomic composition were observed, with mammal species comprising 40.3% of the total species detected during the wet season and 54.2% during the dry season, while the proportions of bird and reptile species were higher in the wet season (Fig. 4). Soil samples had reduced overall species richness, primarily due to fewer bird species (n=11), compared to water (n=41) and sediment (n=37) samples. No amphibian and reptile species were detected in the from soil.

**Fig. 4.**
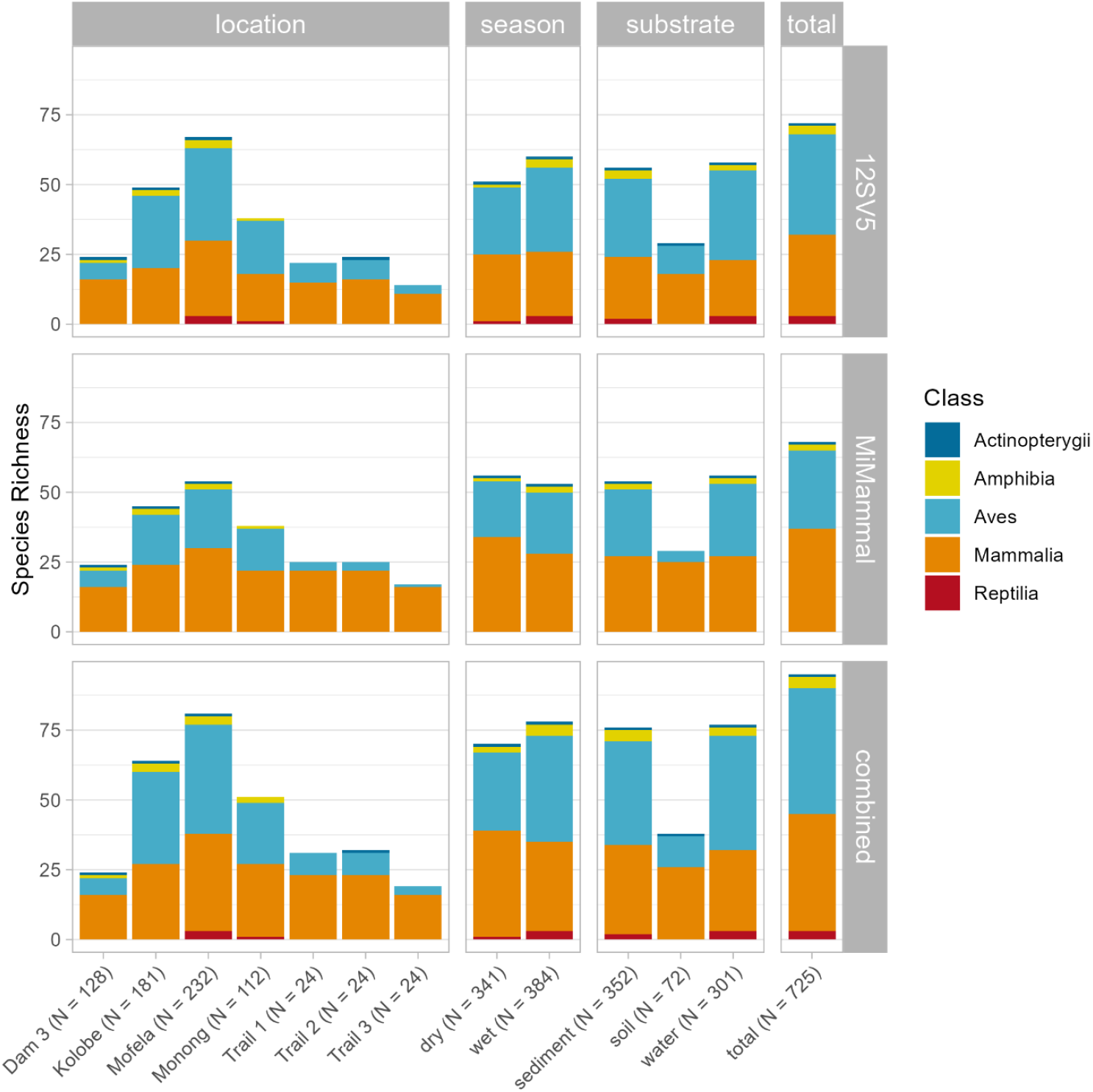
Bar plots showing species richness of each dataset, grouped by location, season, substrate, and overall (total). Results are presented separately for the two primer pairs (12SV5 and MiMammal) and the combined dataset. Colored sections within each bar represent vertebrate classes. Labels on the x-axis give sample sizes in brackets.

The most frequent number of positive samples per species detection in a sampling event was 1 across all substrates, reflecting eDNA’s patchy distribution. Sediment and water showed right-skewed distributions (median: 2), while soil had a more uniform distribution (median: 4; Fig. S4).

NMDS plots and PERMANOVA analysis revealed that species composition was significantly affected by primer pair (F = 110.1, p < 0.001), substrate (F = 25.0, p < 0.001), location (F = 8.3, p < 0.001) and season (F = 7.6, p < 0.001). Time since first sampling day had no significant effect (F = 1.4, p = 0.214). In these analyses, the NMDS space representing Monong and the three trails was largely encompassed within the convex hulls of the three remaining waterholes (Fig. 5a). Water samples covered a much larger area in ordination space compared to sediment and soil samples, which were almost entirely enclosed within the convex hull of the water samples (Fig. 5b). The convex hulls for the dry and wet seasons showed partial overlap with the wet season occupying a larger area in ordination space than the dry season (Fig. 5c). Finally, NMDS plots showed that the regions occupied by the two primer pairs in ordination space were essentially non-overlapping (Fig. 5d).

**Fig. 5.**
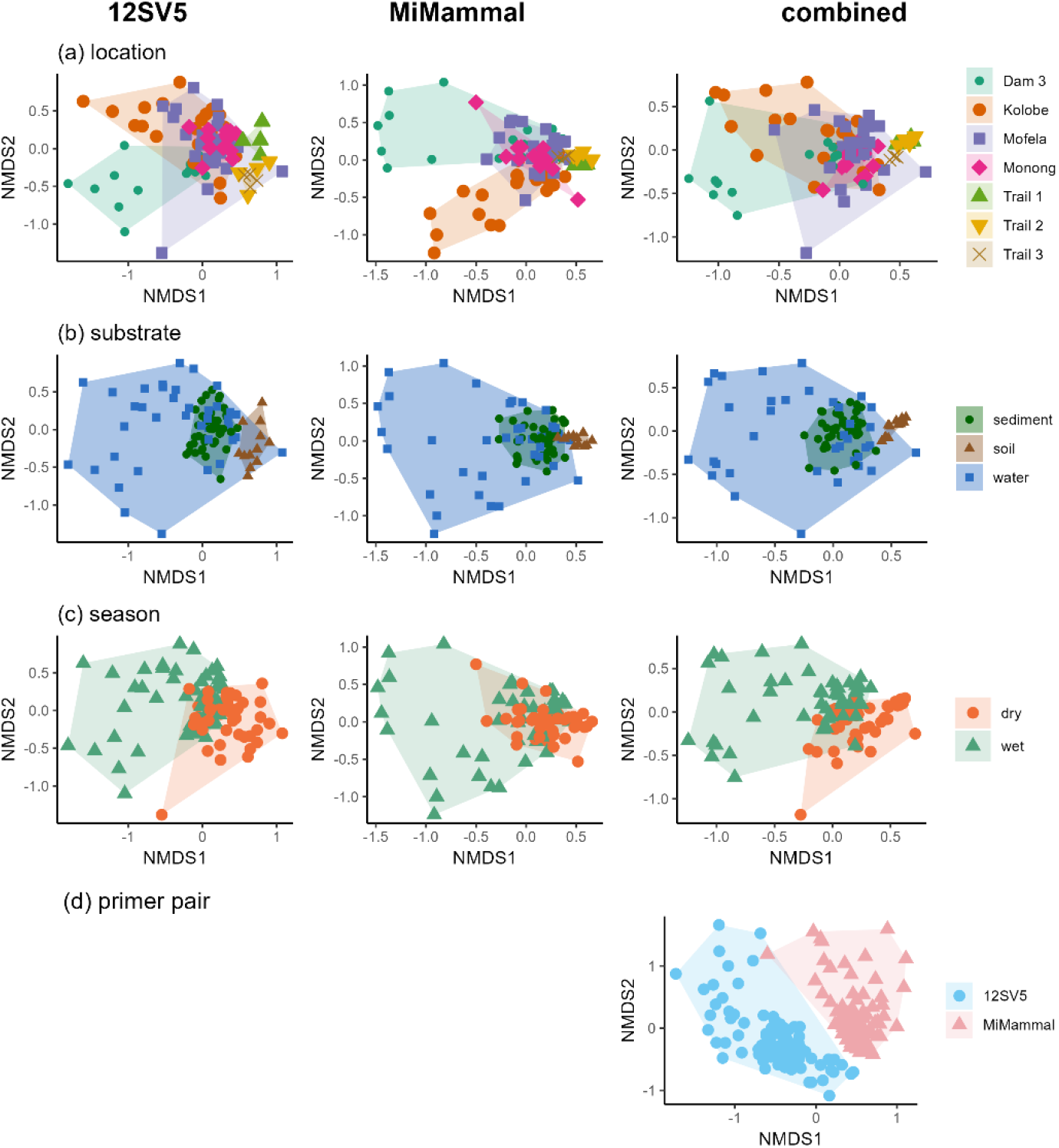
Nonmetric multidimensional scaling (NMDS) plots presenting the differences in species composition among sampled locations (a), substrates (b), between seasons (c) and primer pairs (d). NMDS was performed for the 12SV5 dataset only (column 1), the MiMammal dataset only (column 2) and over both datasets combined (column 3).

While the slope of most species accumulation curves significantly flattened between five and ten sampling events, none of the curves fully levelled off, indicating that new and rarer taxa continued to be detected even after 80 or more sampling events (Fig. 6). Notably, sediment samples detected more species within the first five sampling events, while water samples identified more taxa overall with increased sampling. Both primer pairs detected similar numbers of species, after up to ten sampling events. However, the 12SV5 primer detected more species as sampling efforts increased.

**Fig. 6.**
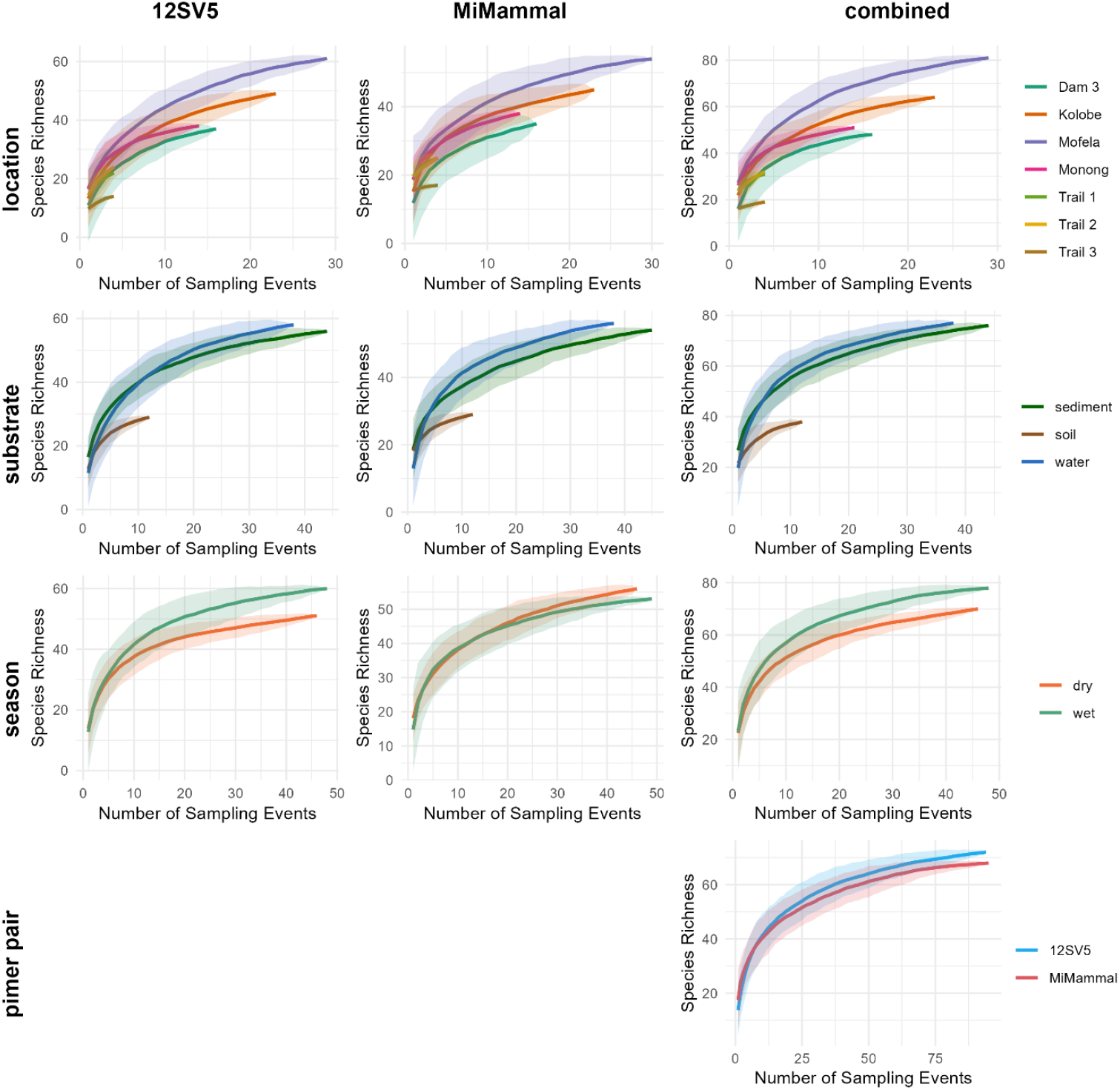
Species accumulation curves showing the number of species detected as a function of the number of sampling events conducted, comparing the sampled locations (a), substrates (b), sampling seasons (c), and primer pairs (d) of this study. Solid lines represent the mean values of species richness, with shaded areas indicating the 95% confidence intervals.

Detection frequencies of species ranged from 0.01 to 0.93 (Fig. 2, Table S8) and did not significantly differ between birds and mammals (W = 828, p = 0.321) but sediment samples resulted in overall higher detection frequencies than water samples (V = 2779, p = 0.001). Sediment samples also yielded higher detection frequencies for abundant species (V = 944, p < 0.001), although this pattern was reversed for rare species (V = 325, p = 0.048). Furthermore, sediment samples yielded significantly higher detection frequencies than water samples for mammals (V = 551, p = 0.003), whereas there was no difference in detection frequencies between sediment and water samples for birds (V = 638, P = 0.096).

The mixed sampling approach resulted in a mean species richness per sampling event similar to sediment-only sampling (26.4 ± 0.9 and 26.8 ± 0.8, respectively, Wilcoxon rank sum test, p > 0.05), but was higher than water-only sampling (19.5 ± 1.3, both p < 0.001, Fig. 7a, Kruskal-Wallis rank sum test; χ^2^ = 22.5, Wilcoxon rank sum tests, both p < 0.001). Overall species richness was higher for the mixed sampling approach (84 species) compared to either single-substrate approach (72 species for sediment and 77 for water, Fig. 7b). The mixed approach consistently yielded higher species richness estimates independent of sampling event numbers, although 95% confidence intervals showed substantial overlap with those of the sediment and water-only approaches (Fig. 7c).

**Fig. 7.**
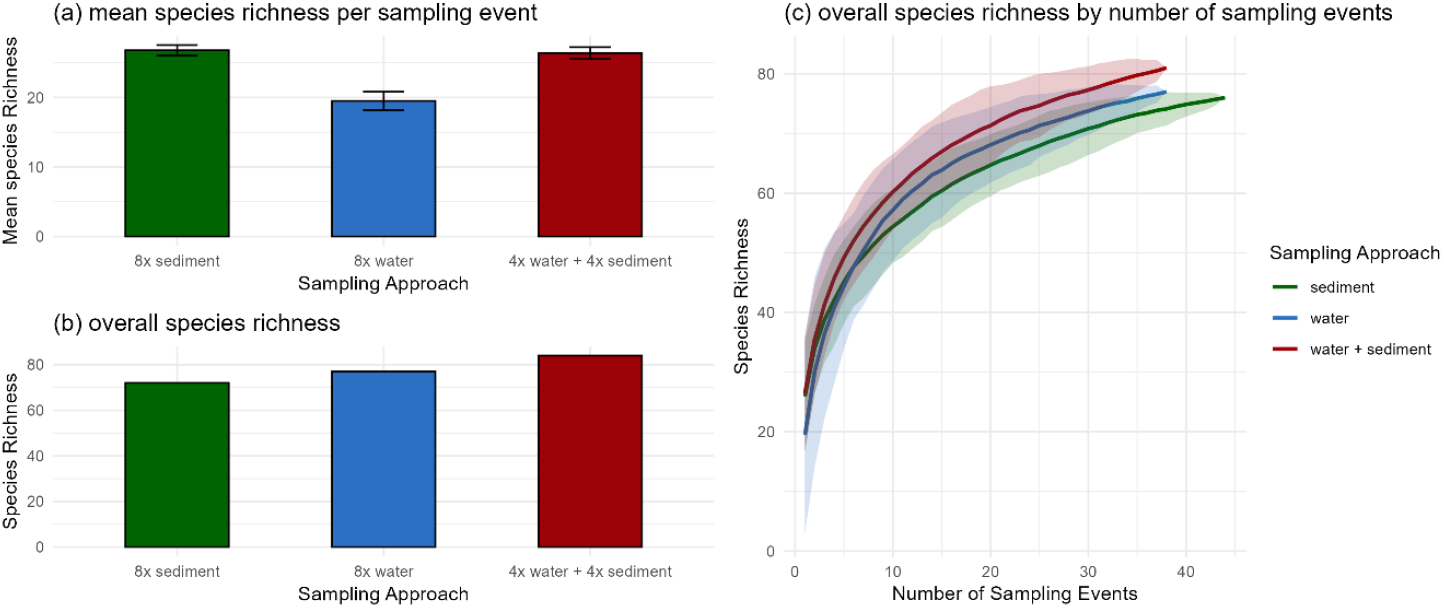
Comparison of the mixed sampling approach (four water samples and four sediment samples per sampling event) to the two single-substrate sampling approaches (eight sediment samples or eight water samples per sampling event). (a) mean species richness per sampling event with error bars showing standard deviation (b) overall species richness over all sampling events (38 for water and mixed, 44 for sediment) (c) species accumulation curves for the three sampling approaches. Shaded areas show 95% confidence intervals

## Discussion

### Overall species richness and potential false positives

The 95 species detected in this study included 19 of the 20 mammal species documented for the game reserve by Morris (2022). The missing mammal was the common reedbuck (*Redunca arundinum*), one of the rarest antelopes in the reserve. Moreover, 23 additional mammals not listed by Morris (2022) were also detected. Of the 100 most common birds recorded for pentad 2530_2540 between 2007 and 2022 (African Bird Atlas Project, 2022), which encompasses most of Botsalano Game Reserve, eDNA metabarcoding detected 25 species, along with 20 additional species not ranked among the top 100. The initial taxonomic assignment revealed a relatively high number of vertebrate species overall without occurrence records in South Africa (39). A small portion of these detections could be attributed to anthropogenic activity but the majority of the remaining cases were very likely due to taxonomic misclassifications, as congeneric documented species frequently lacked reference DNA sequences for the target fragments published in the MIDORI database (Table S3). However, even with the stringent filtering steps we applied, five species are still likely to represent false positives. First, neither the African elephant (*Loxodonta africana*), nor the nyala (*Tragelaphus angasii*), have been introduced to the reserve and the habitat around Botsalano Game Reserve does not meet the species’ requirements, making their presence both within and around the reserve highly unlikely. Neither DNA has ever been processed in any facility involved in the laboratory workflow, excluding contamination during sample processing. No close relatives of the African elephant (Asian elephant, manatees and dugongs) live nearby the sampling area, rendering taxonomic misclassification for this species unlikely. For nyala two congeneric species occur in the reserve, namely the eland (*Tragelaphus oryx*) and the kudu (*Tragelaphus strepsiceros*) but the respective ASVs assigned to the nyala clearly clustered to the two available genetic reference sequences of the nyala as opposed to kudu and eland (data not shown). We hypothesize that the detection of both elephant and nyala DNA originates from secondary DNA input via birds, most likely vultures, which may have acquired the DNA from carcasses or feces in one of the neighboring reserves harboring these species (e.g., Madikwe Game Reserve - 105 km away or Pilanesberg National Park - 130 km away). All samples in which nyala or elephant DNA was detected originated either from Mofela or Monong waterhole, where large groups of vultures were frequently observed during both sampling seasons. Alternatively, the DNA could have been introduced to the reserve by visitor vehicles, potentially adhering to the tires after visits to those game parks. Additionally, three detected bird species are unlikely to occur in or around the game reserve, namely the mallard (*Anas platyrhynchos*), northern pintail (*Anas acuta*) and Hartlaubs’ duck (*Pternonetta harlaubii*). None of these species has been observed during field work. However, two other *Anas* species, the yellow-billed duck (*Anas undulata*) and red-billed teal (*Anas erythrorhyncha*) were frequent visitors of the waterholes and are listed among the top 100 abundant birds for that area (African Bird Atlas Project, 2022), both of which lack genetic reference sequences of both target fragments. We therefore hypothesize that the mallard and pintail detections arose from taxonomic misclassifications from one of these (or other) *Anas* species without genetic reference sequences. Finally, Hartlaub’s duck has only a single recorded occurrence in South Africa in the GBIF database, with its primary distribution range in Central and West Africa. This makes its presence in Botsalano Game Reserve highly unlikely. We attribute the detection of this species to a misclassification of the knob-billed duck (*Sarkidiornis melanotos*), which does occur in the area of the reserve. The 12S rRNA target fragment of these two species differ by only one base pair, which resulted in a slightly better match with Hartlaub’s duck (98/101 bp) than the knob-billed duck (97/101 bp).

Taken together, the potential false positives in our dataset can be attributed to (1) Secondary eDNA input into the study system, which has already been pointed out as an “overlooked source of false positives” for fishes in aquatic urban systems (Xiong et al., 2024), but has not yet been investigated thoroughly in terrestrial systems; (2) Incomplete genetic reference databases, which is a persisting topic in eDNA metabarcoding studies, particularly for understudied taxa (Keck et al., 2023; Schenekar et al., 2020), and (3) low genetic resolution of metabarcoding markers, which is a concern particularly for young and/or sister species and would require the utilization of other or additional metabarcoding markers (Keck et al., 2023). All three of them underscore the importance of taxonomic and ecological expertise in critically evaluating eDNA metabarcoding results.

### Effect of workflow variations Sampled substrate

Water and sediment samples showed similar overall species richness and taxonomic compositions, though with differences in detection patterns regarding species abundance. Sediment samples yielded higher detection frequencies for abundant species across sampling events, whereas water samples were more effective at detecting rare taxa (Fig. 2). Furthermore, sediment samples exhibited higher species richness per sampling event (Fig. 7a) and displayed a flatter decline of number of positive samples per species detection (Fig. S4), indicating the more consistent detection of abundant taxa across samples and sampling events. In contrast, water samples yielded a greater overall species richness across the same number of sampling events (Fig. 7b) and a larger NMDS space occupied (Fig. 5b), both of which reflect a higher number of rare, exclusive species detections from water samples but also higher variability in species composition across sampling events. This pattern may result from the longer eDNA persistence in sediment (Corinaldesi et al., 2005; Sedlmayr & Schenekar, 2024; Turner et al., 2015) and the higher concentration of abundant taxa eDNA along shorelines, which may outcompete rare taxa in metabarcoding. The larger water sample volumes (compared to sediment) may have counteracted eDNA patchiness and increased the detection of rare DNA fragments from low-abundance species. (Altermatt et al., 2023; Sepulveda et al., 2019). Sediment sampling was quicker and required minimal equipment—gloves, plastic spatulas, and sampling tubes—with limited bleaching. In contrast, filtering a single water sample took up to 25 minutes, including pre-filtering, filtration, and equipment handling. Sediment samples were processed in under two minutes by adding Longmire buffer, sealing, and shaking, making them significantly more field-efficient.

Soil samples collected from wildlife trails around waterholes yielded a lower species richness overall but were collected at a much lower sampling numbers than water and sediment samples. However, mammalian species richness was comparable to that detected from water and sediment samples. The reduced overall species richness primarily resulted from fewer bird detections. While terrestrial mammals frequently use these trails to access waterholes, depositing eDNA in the soil, bird eDNA is more abundant directly at the waterhole, where eDNA deposition results from direct interactions with the water, such as drinking, or defecating while standing in water or being perched on overhanging branches. Thus, wildlife trails around waterholes may serve as viable alternative sampling sites for monitoring terrestrial mammals, particularly when direct sampling at waterholes is impractical due to inaccessibility or the presence of dangerous animals at the time of sampling.

### Seasonality

Overall, we detected more vertebrate species in the wet season than the dry season but more mammals were detected during the dry season. While differences in filter types used for sampling between seasons cannot be ruled out as a contributing factor, we primarily attribute this pattern to two main ecological factors: 1) The increased biological activity of birds, amphibians and reptiles during the wet season, which coincides with the peak reproductive period of most species (Moreau, 1950), led to higher detection probabilities for these taxa. 2) For non-migratory mammals that also do not hibernate through the dry season, the reduced number of water bodies in the reserve (only two waterholes remained at the end of the dry sampling season) led to a higher utilization of the sampled waterholes, increasing detection probabilities and overall detected mammalian species richness.

### Primer pair

The two primer pairs tested had the strongest influence on the detected species composition, with virtually non-overlapping NMDS areas occupied (Fig. 5d) and only 47% of species overlap (Fig. 3). The vertebrate-specific 12SV5 primer pair captured a broader range of vertebrate diversity, whereas the mammal-specific MiMammal primer pair identified more mammal species. However, notably the 12SV5 primer detected five mammal species that the MiMammal primer pair did not while the MiMammal primer pair detected 31 non-mammal vertebrates, including 10 undetected by 12SV5, highlighting its capacity to detect taxa beyond its primary target group. Consistent with previous studies (McElroy et al., 2020; Wang et al., 2023), our results also support the use of at least two primer pairs to maximize species detection in eDNA metabarcoding studies.

## Conclusions and recommendations

The findings of this study provide guidance for optimizing eDNA metabarcoding sampling designs around waterholes in semi-arid or arid systems, like savannas, targeting terrestrial vertebrates. Regarding substrate choice, sediment samples performed superior in the consistent detection of abundant species, independent of taxonomic class (e.g., birds or mammals), while water samples were more effective for detecting rare species. Therefore, for studies aiming to broadly assess species composition in a new or poorly studied system—or where field conditions limit sample processing logistics, such as remote locations or extensive daily sampling—we recommend sediment sampling of water bodies due to its simplicity and efficiency. Conversely, if detecting rare taxa is a priority and field conditions permit timely water filtration, water samples should be preferred. A combined sampling approach incorporating both substrates is likely to yield the highest species richness.

Soil samples from wildlife trails near waterholes offer a promising alternative for monitoring terrestrial mammals when direct access to waterholes is not possible. However, they are less effective for detecting other taxa, such as birds, amphibians, and reptiles, which rely more on direct interactions with water for eDNA deposition.

Sampling during the dry season may enhance terrestrial mammal detection, as these species tend to concentrate around the limited remaining water sources. In contrast, the detection of other taxa may decrease due to lower biological activity or migration. Additionally, seasonal changes in physicochemical properties—particularly suspended particle load—can significantly impact sampling efficiency, necessitating workflow adjustments, such as selecting appropriate filter pore sizes for water filtration. Using two primer pairs had the greatest positive impact on detected species richness. To maximize species detection within a given budget, we recommend prioritizing two independent genetic markers— or more where feasible—over e.g., adding an extra sampling season.

### Data archiving, funding & benefit sharing statement

The sequencing data of this study have been deposited in the GenBank Sequence Read Archive (SRA) under accession number PRJNA1218494 and can be accessed at https://www.ncbi.nlm.nih.gov/sra/PRJNA1218494. Benefits generated: A research collaboration was developed with scientists from the Austria and South Africa and all scientists actively contributing to logistic organization, data generation, data analysis and manuscript writing are either listed as co-authors or mentioned in the acknowledgements. The results of this collaborative project are being frequently shared with all collaborators and the broader scientific community. The research addresses a priority research focus of our South African collaborator (SANBI), namely biodiversity monitoring in South Africa. Sampling for the project has been approved by North West Parks Board and by the Department of Economic Development, Environment, Conservation and Tourism, South Africa (Permit No. NW 39881/05/2022), the research has been approved by SANBIs NZG Animal Research Ethics and Scientific Committee (P2022/15) and a permission in terms of section 20 of the animal diseases act 1984 was received from the Department of Agriculture, Land Reform and Rural Development, South Africa (DALRRD). A certificate of compliance in accordance with Article 17, paragraph 2, of the Nagoya Protocol has been issued by the Access and Benefit-Sharing Clearing-House (ABSCH) of the Department of Forestry, Fisheries and the Environment, South Africa (ABSCH-IRCC-ZA-262932-1). This research was funded in whole, or in part, by the Austrian Science Fund (FWF) [10.55776/P35059]. For the purpose of open access, the author has applied a CC BY public copyright license to any Author Accepted Manuscript version arising from this submission.

## Supplementary material

**Supplementary material 1.docx:** Contains Figs. S1, S2, S3 & S4 and Table S1

**Table S3.xlsx** contains Table S3

**Table S4.xlsx** contains Table S4

**Table S5.xlsx** contains Table S5

**Table S6.xlsx** contains Table S6

**Table S7.xlsx** contains Table S7

**Table S8.xlsx** contains Table S8

