## Supplementary material 1 for "Unveiling vertebrate biodiversity in arid and semi-arid terrestrial ecosystems through eDNA metabarcoding at savanna waterholes"

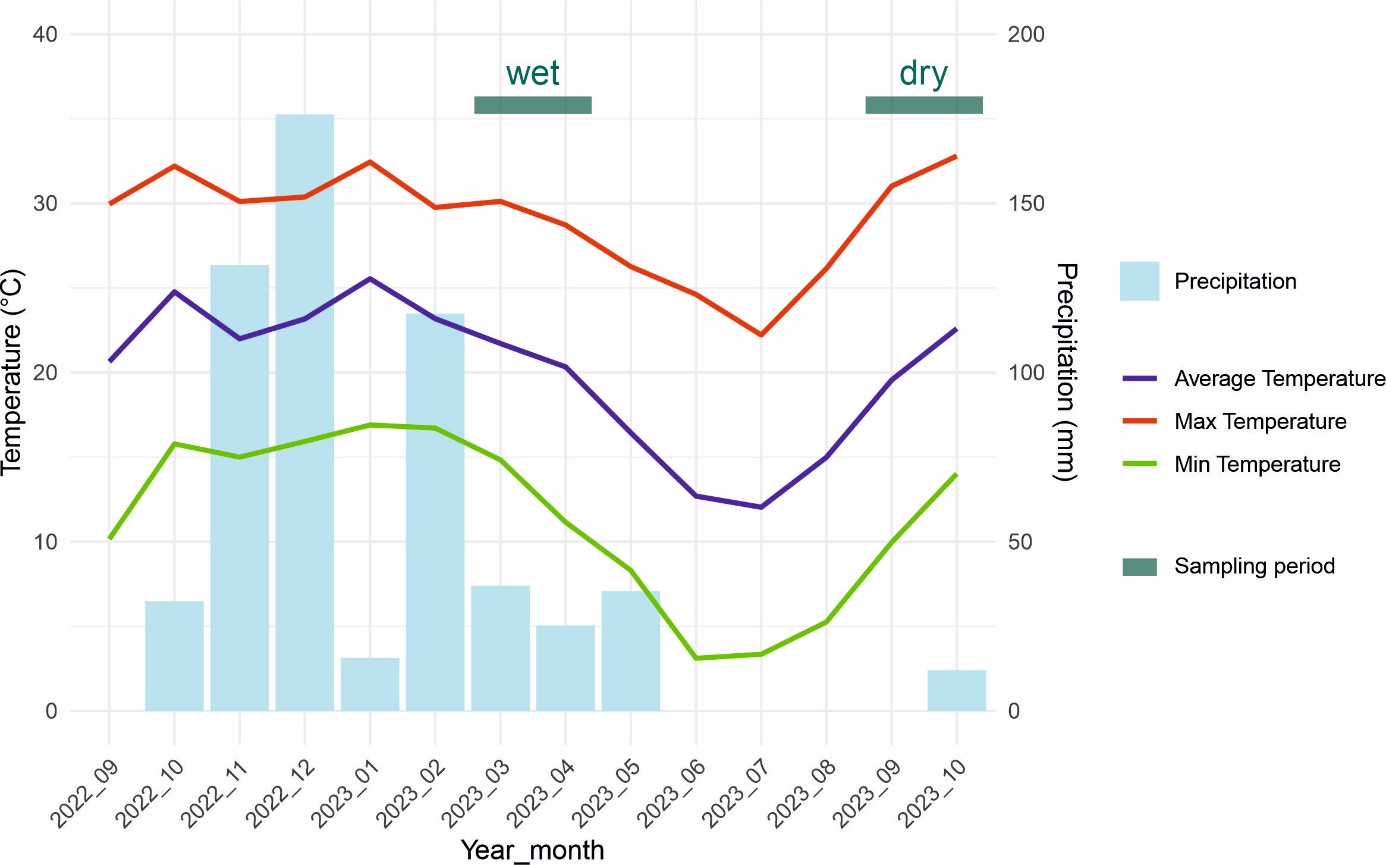

**Fig. S1** Average, maximum and minimum daily temperature, as well as average monthly rainfall (precipitation) of the study area in the time frame of the fieldwork. The sampling periods of the two seasons (“wet” and “dry”). are indicated by the two green bars. Precipitation data was received from Botsalano Game Reserve management, and temperature data from the National Centers for Environmental Information / National Oceanic and Atmospheric Administration (NCEI/NOAA).

**Table S1** Larger-bodied mammals recorded in Botsalano game reserve in October 2020. From (Morris, 2022). This count did not target small animals such as rodents or bats.

| **Common name** | **Scientifc name** | **Tswana name** | **Count** |
| --- | --- | --- | --- |
| Black wildebeest | *Connochaetes gnou* | Kgokong/*Pudumô | 374 |
| Black-backed jackal | *Canis mesomelas* | Phokojê | 10 |
| Blesbok | *Damaliscus pygargus phillipsi* | *Nônê | 343 |
| Brown hyena | *Parahyaena brunnea* | Phiri | 1 |
| Buffalo | *Syncerus caffer* | *Nare | 25 |
| Burchell's zebra | *Equus quagga* | *Pitse ee Tilodi | 168 |
| Common duiker | *Sylvicapra grimmia* | Photi | 9 |
| Common reedbuck | *Redunca arundinum* | Mofele oo Mosetiha | 2 |
| Eland | *Tragelaphus oryx* | *Phôfu | 3 |
| Gemsbok | *Oryx gazella* | *Kukama | 212 |
| Giraffe | *Giraffa giraffa* | *Thutlwa | 34 |
| Impala | *Aepyceros melampus* | *Phala | 268 |
| Kudu | *Tragelaphus strepsiceros* | *Thôlô | 57 |
| Mountain reedbuck | *Redunca fulvorufula* | Mofele o mohibidu | 3 |
| Red hartebeest | *Alcelaphus buselaphus* | *Kgama | 161 |
| Springbok | *Antidorcas marsupialis* | Tshêpê | 439 |
| Steenbok | *Raphicerus campestris* | Phuduhudu | 8 |
| Vervet monkey | *Chlorocebus pygerythrus* | Kgabo | Not counted |
| Warthog | *Phacochoerus africanus* | Makôrwane/*Kolobe | 158 |
| Waterbuck | *Kobus ellipsiprymnus* | *Motomoga | 44 |
| White rhino | *Ceratotherium simum* | Tshukudu | Confidential |

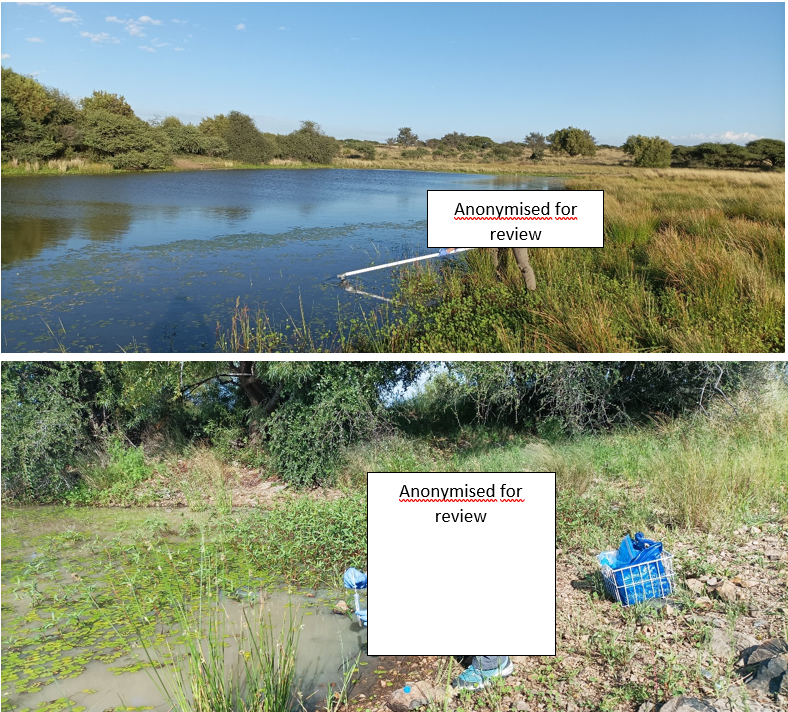

**Fig. S2** Water (a) and sediment (b) sampling at the waterholes.

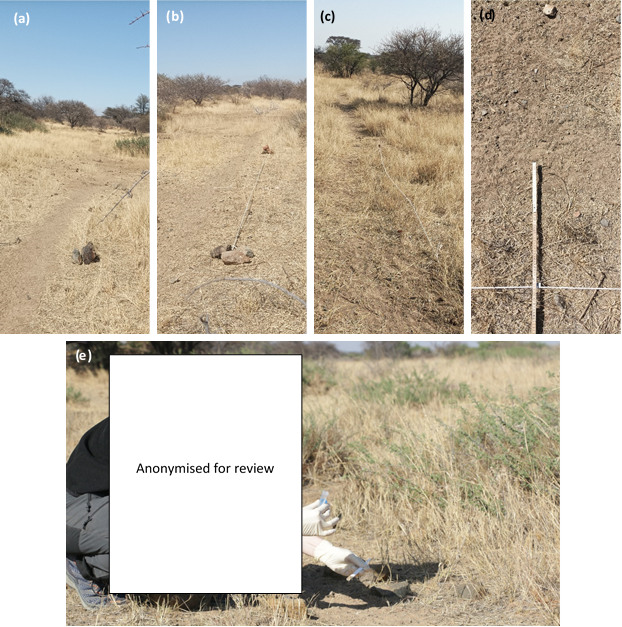
**Fig. S3** Wildlife trail transects for soil sampling. Layout of Trails 1, 2, and 3 (a-c). For each transect, six sampling points were marked at 1 m intervals on the rope, and sampling was conducted 50 cm into the wildlife trail from the transect line (d) using a DNA-free spatula and 15 tube (e).

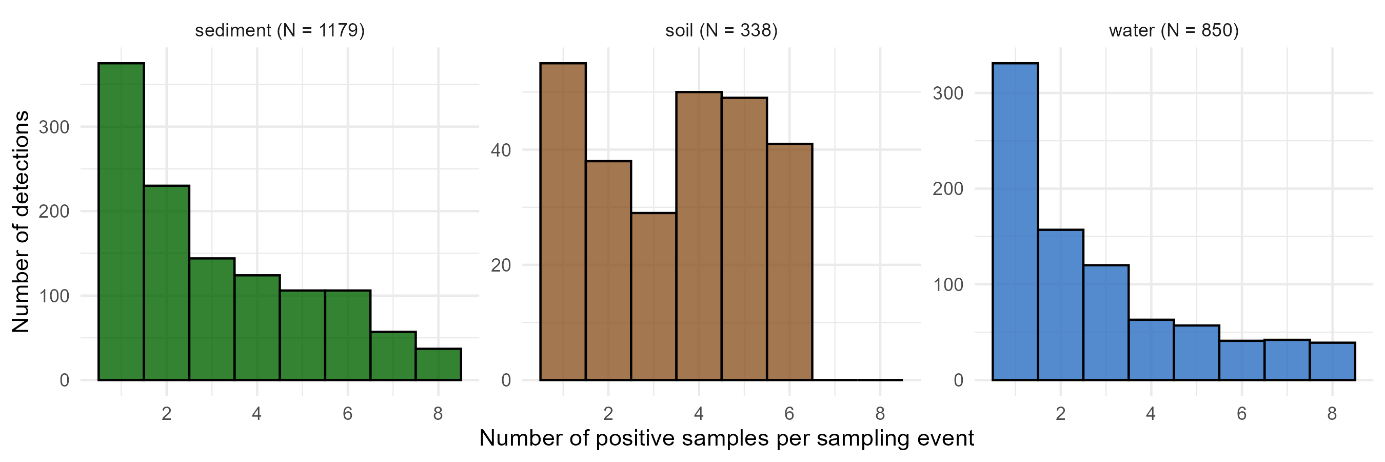
**Fig. S4** Histograms showing the frequencies of the number of positive samples once a species has been detected in a sampling event. Per sampling event, eight samples were collected for sediment and water samples, whereas for soil samples, six sampling replicates were collected.
